# A metastable Gα_S_/β-arrestin complex sustains endosomal PTH1R signaling

**DOI:** 10.64898/2026.11.11.765718

**Authors:** Jonathan Pacheco, Satyaki Saha, Karina A. Peña, Ivet Bahar, Jean-Pierre Vilardaga

**Author notes:** Correspondence. Email: I.B.; J.-P.V. These authors contributed equally to this work (Jonathan Pacheco, Satyaki Saha, Karina A. Peña).

## Abstract

G protein-coupled receptors (GPCRs) initiate cellular signaling by activating heterotrimeric G proteins at the plasma membrane and are silenced when β-arrestin (βarr) binds the phosphorylated receptor, preventing G protein binding. However, many receptors continue to generate cAMP after internalization, even while βarr is bound, a paradox that lacked a molecular mechanism in live cells. A direct β-arrestin1–Gαs interface in the PTH1R-βarr1-Gαs complex has recently been defined structurally, but whether the complex operates in endosomes, and how it behaves there, has not been established. Using intact cells, we examine the assembly at three spatial resolutions. Confocal imaging with a nanobody selective for nucleotide-free Gαs places activated Gαs, βarr, and agonist-occupied PTH1R on the same endosomes. Single-molecule localization microscopy, with illumination modes that separate the plasma membrane from the endosomal membrane in single cells, shows that the agonist recruits Gαs and βarr at both membranes. FRET, however, reveals a specific interaction only on endosomes, where a Gαs inhibitor restricted to that compartment abolishes the sustained cAMP phase without affecting receptor internalization. Weighted-ensemble-enhanced molecular simulations show that the complex is metastable, releasing Gαs through a large rigid-body rotation away from βarr while βarr itself tightens its interactions with the receptor. These findings provide a molecular basis for endosomal Gs/cAMP signaling and open a new avenue for drug development targeting this pathway.

**One Sentence Summary:** Gα_S_/β-arrestin/PTH receptor complex as endosomal signaling intermediate.

**Significance Statement:** Hormone receptors were long thought to be switched off once β-arrestin binds and relocates them inside the cell. However, several receptors keep signaling by producing cAMP from endosomes, and how they do so is unclear. Using live cells, we show that the parathyroid hormone receptor, β-arrestin, and the Gαs subunit form a complex on endosomes but not at the cell membrane, even though all three accumulate at both membranes. Blocking Gαs only on endosomes removes the sustained phase of the hormone response without affecting receptor endocytosis. Molecular simulations show the complex is unstable, yet detectable in cells: Gαs leaves the complex by a large rotation that requires the receptor to open further, while β-arrestin holds the receptor more tightly, allowing repeated rounds of activation. β-arrestin therefore plays a critical role in sustaining endosomal signaling.

## Introduction

GPCRs transduce extracellular signals by coupling to heterotrimeric G proteins at the plasma membrane, promoting GDP–GTP exchange on the Gα subunit (1–3). The resulting dissociation of GTP-bound Gα allows for the engagement of both Gα and Gβγ with downstream effectors; for stimulatory Gα_S_, this activates transmembrane adenylyl cyclases and increases cyclic AMP (cAMP) production (4). Signaling is then terminated by a two-step mechanism: G protein-coupled receptor kinases (GRKs) phosphorylate the agonist-bound receptor on its intracellular loops and C-terminal tail, and βarr binds the phosphorylated receptor, sterically occluding the intracellular cavity otherwise recognized by G proteins while simultaneously engaging the receptor in dynamin-dependent internalization (5–8). In this canonical framework, βarr and G proteins are functionally antagonistic, because βarr recruitment terminates receptor/G protein coupling and signaling (7, 9). However, accumulating evidence across multiple receptors shows that G protein/cAMP signaling continues after receptor internalization and βarr engagement (10–13). This sustained endosomal signaling produces pharmacological responses that are temporally and spatially distinct from those arising at the plasma membrane (14, 15), and the classical desensitization model cannot explain this process. How internalized, βarr-bound receptors continue to activate G proteins remains a central unresolved question in GPCR biology.

The parathyroid hormone (PTH) receptor type 1 (PTH1R) serves as a paradigm for this problem. PTH stimulation initiates a first acute wave of cAMP production at the plasma membrane through canonical Gs coupling. βarr binding mediates the internalization of PTH-bound PTH1R to early endosomes, where the receptor generates a second, sustained wave of cAMP that is pharmacologically distinct from the first and required for calcium homeostasis and vitamin D activation (reviewed in (16)). Receptor recycling via the retromer complex subsequently terminates this endosomal phase and resets the system for another signaling cycle (17). The recognition of these cAMP-signaling waves, now documented for multiple Gs-coupled GPCRs, including class A and B GPCRs such as GLP-1R, GCGR, and V2R (see recent review (13)), deepens the existing paradox. If βarr blocks G protein access to the receptor’s intracellular core, how does the internalized, βarr-bound receptor continue to activate Gs from the endosomes?sssssss

A partial answer came initially from the demonstration that a single receptor can engage G protein and βarr simultaneously. These complexes, termed megaplexes, form preferentially at receptors whose phosphorylated C-terminal tails bind βarr strongly, and they sustain G protein signaling from internalized compartments (18). The cryo-EM structure of an endosomal β_2_-adrenergic/V2R chimera–Gs–βarr1 megacomplex subsequently showed that all three components can adopt their canonical active conformations at once, with the G protein engaging the receptor core and βarr the phosphorylated tail (19), and comparable assemblies have since been visualized for other receptors. Prior to these studies, we showed that βarr can prolong and sustain endosomal PTH1R signaling via cAMP by increasing the steady-state level of activated Gs (20), presumably enabling multiple rounds of Gαs activation after internalization. Whether a direct endosomal βarr–Gαs interaction contributes to it remains unknown.

Recent structural studies show that βarr1 interacts with PTH1R either via its finger loop inserted into the receptor core (21), a configuration that precludes G protein binding, or through the phosphorylated C-terminal tail alone, which leaves the intracellular cavity of the transmembrane domain (TMD) accessible, as initially observed in the glucagon receptor (GCGR)–βarr1 complex (22, 23). This structural flexibility has led to the hypothesis that the core-engaged state mediates desensitization and internalization at the plasma membrane, whereas the phosphorylated tail-engaged state in endosomes allows reassembly with diffusing Gαs, resulting in sustained signaling. However, this hypothesis is inconsistent with the ability of the phosphorylation-deficient PTH1R to interact with βarr, internalize, and maintain sustained cAMP (24). It also remains unresolved whether a PTH1R–Gαs–βarr assembly forms in the endosomal compartment and evolves over time.

Here we address this question in intact cells and *in silico*. Using TIRF and HILO to optically separate the plasma membrane from endosomes, endosomal FRET and conformational-nanobody imaging restricted to endosomes, and an endosome-targeted Gα_s_ inhibitor, we show that PTH1R activation induces formation of a βarr1-Gαs complex selectively on endosomes. We then perform weighted-ensemble (WE)-enhanced molecular dynamics (WE-MD) simulations as well as conventional microseconds-long MD simulations, initiated from the cryo-EM structure of the PTH1R–βarr1–Gαs megaplex determined independently and reported separately (25) to show that the complex is metastable and that Gα exhibits a tendency to dissociate from the megaplex while βarr remains bound to the receptor and even further tightens its interactions with the receptor. Gα_S_ destabilization/dissociation, accompanied by stronger PTH1R-βarr1 association, is enhanced in the presence of GTP-bound Gs. The simulations of the megaplex dynamics provide first insights into the time-resolved antagonistic effect of receptor-βarr1 association on receptor-Gαs interactions.

## Results

### Nucleotide-free Gαs and β-arrestin are recruited to the same endosomes

Our previous work showed that β-arrestins are required for the prolonged cAMP response generated by PTH-activated PTH1R in early endosomes and proposed that they act by maintaining the steady-state level of activated Gs (17, 20). To account for this observation, we tested the hypothesis that a ternary PTH1R–Gα_S_–βarr complex (hereafter the PTH1R megaplex) assembles on endosomes to prolong receptor signaling via cAMP.

We examined whether PTH1R, Gαs, and βarr occupy a common endosomal assembly at three successively higher spatial resolutions, each addressing a different aspect of the question: confocal microscopy, which resolves ∼250 nm and therefore reports whether the components reach the same endosomes; nanoscale localization microscopy, which resolves ∼100 nm and reports how they are distributed within that organelle; and FRET, which is sensitive over ∼10 nm and reports the direct protein-protein interactions themselves.

At the lowest resolution, we asked whether activated Gαs is present in endosomes containing βarr. To this end, we exploited the selectivity of the nanobody Nb37 for nucleotide-free Gαs as a specific marker for the active state. Confocal microscopy showed that PTH induced endosomal colocalization of βarr1 fused to monomeric Tomato (βarr1^Tom^) with Nb37 fused to GFP (Nb37^GFP^) (**Fig. 1A**), also colocalized with TMR-labeled PTH (PTH^TMR^) in endosomes (**Fig. 1A**, *arrows*). Before stimulation, Nb37^GFP^ was distributed diffusely, and no Nb37-positive vesicles were detected. LA-PTH stimulation produced discrete Nb37^GFP^-positive endosomes and within those objects βarr1^Tom^ and Nb37^GFP^ signals corresponded closely, with a mean per-cell Pearson’s correlation coefficient of 0.76 ± 0.13 [mean ± SD] (**Fig. 1B**). The agonist dependence of the recruitment is given by its time course: endosomal Nb37^GFP^–βarr1^Tom^ colocalization was absent before stimulation and developed over 10 minutes after ligand addition (*SI Appendix* **Fig. S1**), showing that the Nb37-positive endosomal pool is generated by receptor activation. At the resolution of confocal microscopy, these images show that nucleotide-free Gα_S_ and βarr1 co-localized to the same endosomes in response to PTH1R activation. However, they cannot show how the proteins are distributed within the membrane.

**Figure 1.**
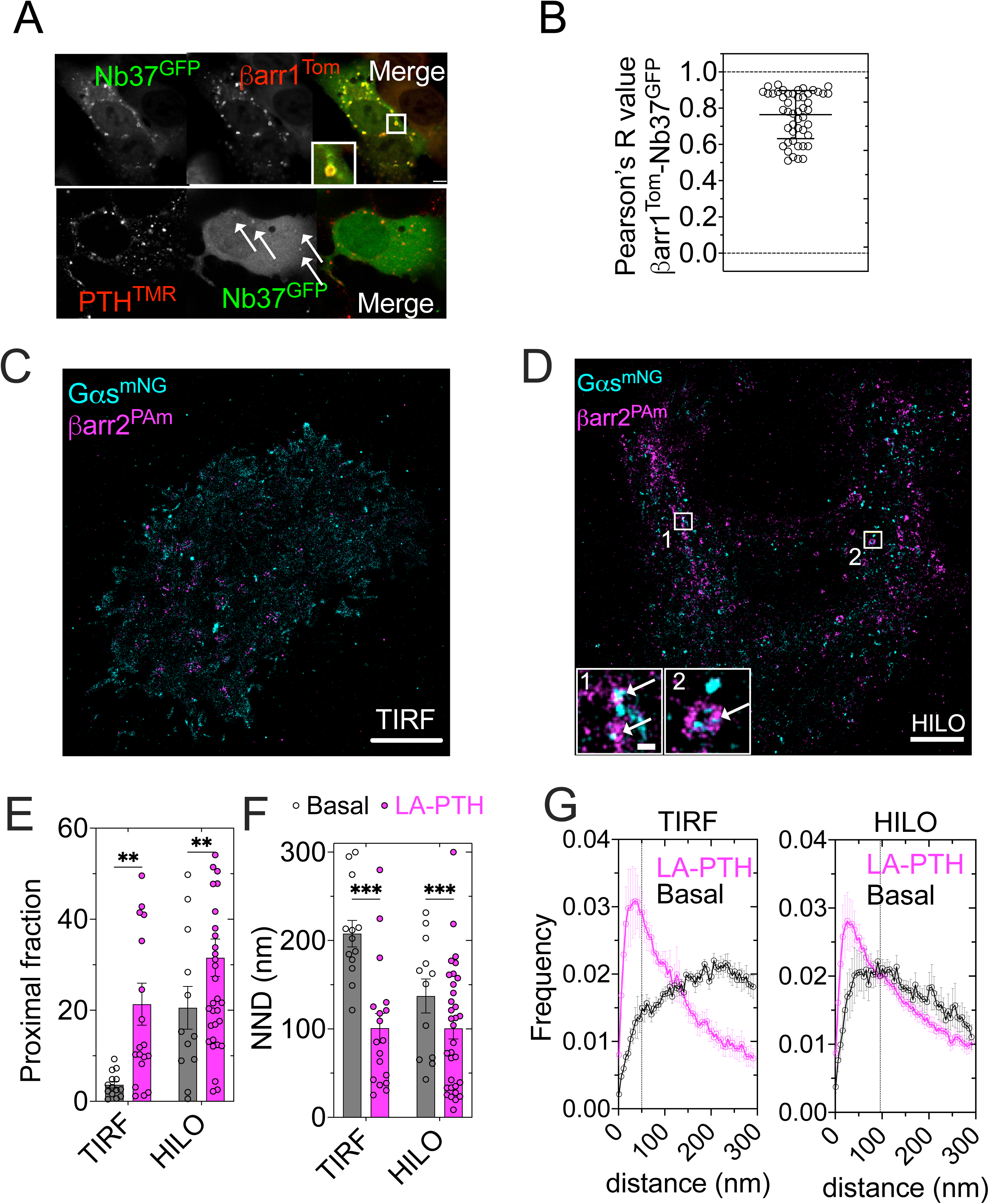
Nanoscale organization of Gαs and β-arrestin at the plasma membrane and on endosomes. **(A, B)** Confocal imaging of Nb37^GFP^ and βarr1^Tom^ distribution in HEK293-R cells stimulated with PTH (top panels) or PTH^TMR^ (bottom panels). Representative microfilms show distribution 15 min post-stimulation. Full time courses are shown in Figure S1. Scale bar, 10 μm. **(C, D)** Representative two-color super-resolution image under TIRF (C) and HILO (D) mode from HEK293 cells coexpressing PTH1R, Gα_S_^mNG^ (*cyan*) and βarr2^PAmCherry^ (*magenta*) after LA-PTH stimulation. Insets 1 and 2 show the boxed regions enlarged, with βarr2 arranged in a ring-like structure of endosomal dimensions and Gα_S_. Scale bar, 5 µm. **(E)** Fraction of Gα_S_^mNG^ with βarr2^PAmCherry^ localizations within 46-94 nm under basal (*open symbols, gray bar*) and after LA-PTH stimulation (*closed symbols, magenta bar*) under HILO illumination. Mean ± sem of n = 10 (TIRF and HILO, basal), n = 14 (TIRF + LAPTH) and n = 25 cells (HILO + LAPTH) from N = 3 independent experiments, with \**P* < 0.05, \*\*\**P* < 0.01, \*\*\*\**P* < 0.001 by two-way ANOVA with a Fisher’s LSD test for multiple comparisons. **(F, G)** Frequency distributions of Gα_S_^mNG^ to βarr2^PAmCherry^ NNDs in TIRF (F) and HILO (G). Data are the mean ± sem. The vertical dashed line indicates the estimated spatial resolution of the experimental setup (46 nm and 94 nm for TIRF and HILO, respectively).

### Agonist-dependent redistribution of Gαs and β-arrestin at the plasma membrane and on endosomes at nanoscale resolution

We imaged Gαs fused to monomeric NeonGreen (Gαs^mNG^) and βarr2 fused to photoactivable mCherry (βarr2^PAm^) in cells by two-color single-molecule localization microscopy, using two illumination approaches that differentiate the plasma membrane (PM) from the endosomal membrane compartments (*SI Appendix* **Fig. S2A**): total internal reflection fluorescence (TIRF), in which the evanescent field extends ∼100 nm from the coverslip and excites the plasma membrane while leaving endosomal vesicles above it dimly excited, and highly inclined and laminated optical sheet (HILO) illumination, in which the refracted light sheet crosses the cytoplasm and illuminates endosomes.

To verify that we can selectively optically separate the plasma membrane from endosomes, we calibrated the laser angle using cells co-expressing the GFP-labeled Early Endosome Antigen 1 (EEA1^GFP^) as an endosomal marker and iRFP-fused Lyn11 (Lyn11^iRFP^) as a plasma membrane marker (**Fig. S2B**). In TIRF mode, the Lyn11^iRFP^ signal was distributed homogeneously, consistent with its PM location, whereas the EEA1^GFP^ signal was dim and diffuse. In HILO mode, Lyn11^iRFP^ appeared as a thin membrane border outlining the cell periphery, whereas EEA1^GFP^ was distributed in multiple discrete endosomes (**Fig. S2B** *inset*). The spatial resolution of each illumination mode was determined by Fourier ring correlation at a fixed 1/7^th^ threshold (**Fig. S2C-D**), yielding 46 ± 0.002 nm for TIRF and 94 ± 0.004 nm for HILO (*n* = 3 and 4, respectively).

Under HILO illumination, βarr2^PAm^ localizations were arranged in ring-like structures of the dimensions expected for early endosomes, with Gαs^mNG^ localized to these rings (**Fig. 1D**). We observed coincident events on endosomes between Gαs and βarr (n per cell) in HILO reconstructions that were absent from TIRF reconstructions (**Fig. 1C, D**). To quantify the spatial relationship between the two proteins, we computed, for each Gαs localization, the distance to the nearest βarr localization, and summarized these nearest-neighbor distances (NNDs) in three ways: as the fraction of Gαs localizations with a βarr neighbor within a proximity threshold defined by the effective spatial resolution, 46 and 94 nm for TIRF and HILO, respectively (**Fig. 1E**) as the mean NND per cell (**Fig. 1F**), and as the full NND frequency distribution out to 300 nm (**Fig. 1G**).

The TIRF assay serves as a positive control for this analysis. β-arrestin recruitment to the plasma membrane upon agonist stimulation is a well-established event in GPCR pharmacology, so a quantification that failed to detect increased Gαs–βarr proximity at the cell surface after LA-PTH would be questionable. At the plasma membrane, LA-PTH increased the proximal fraction from ∼5% to ∼20% and shortened the median NND from ∼200 nm to ∼100 nm (**Fig. 1F, G**). The full distribution shifted from a broad profile toward 150–300 nm to one dominated by a peak below 100 nm, with a clear depletion of long distances (**Fig. 1G**, left). On endosomes, the basal distribution favored shorter distances compared to the plasma membrane (averaged NND ∼140 nm versus 200 nm). LA-PTH induced a smaller but statistically significant change, increasing the proximal fraction from ∼20% to ∼31% and reducing the averaged NND to ∼100 nm (**Fig. 1G**, *right*). Receptor activation therefore induced Gαs and βarr recruitment into both plasma and endosomal membranes. The TIRF assay provided the expected prediction, thereby validating what the NND measurements can report. At the plasma membrane, the agonist effect is large, as expected from β-arrestin recruitment to the plasma membrane, where Gαs is already present. Yet the complementary assays described in the next paragraph find no direct interaction at the PM: single-molecule imaging detected no Gαs–βarr collision events at the cell membrane, and we obtained no evidence of a specific Gαs–βarr FRET signal. Increased proximity at these distances is therefore not sufficient to demonstrate complex formation, which is unsurprising given that a NND of ∼100 nm is an order of magnitude greater than the dimension of a Gα_S_–βarr1 complex. However, the analysis shows that receptor activation co-recruits Gαs and βarr into nanoscale domains of single endosomes.

### Agonist-induced Gαs–βarr interaction on endosomal membranes

To determine whether Gαs and βarr are in direct contact on plasma and endosomal membranes, FRET was measured via acceptor photobleaching in cells co-expressing Gαs^mNG^ and βarr1^Tom^ and stimulated by LA-PTH (**Fig. 2A, B**). Photodestruction of the acceptor in an endosomal region of interest resulted in the expected reciprocal change in donor and acceptor intensities (**Fig. 2B**). To distinguish a specific interaction from proximity-dependent energy transfer between Gαs^mNG^ and βarr1^Tom^, we examined how FRET efficiency (E_app_) depends on the acceptor-to-donor fluorescence ratio (F_Tom_/F_NG_), which rises linearly with acceptor density for non-specific transfer but saturates when donors are engaged specifically in a complex.

**Figure 2.**
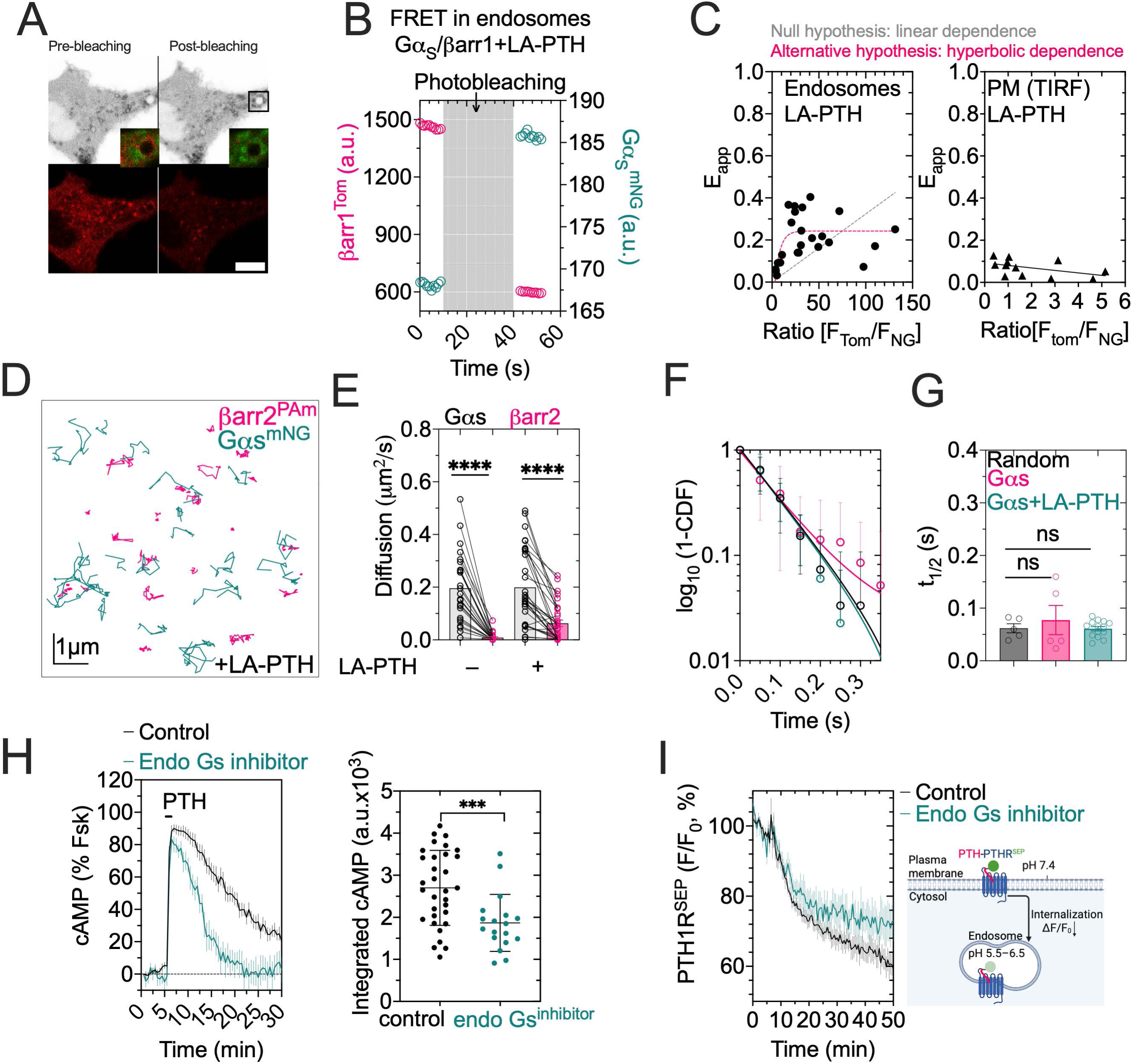
Endosomal PTH1R–Gα_s_–βarr1 complex containing nucleotide-free Gα_s_ sustains cAMP production. **(A, B)** Representative photobleaching experiment in endosomes of HEK293-R cells expressing Gαs^mNG^ and βarr1^Tom^ (A). Effect of photobleaching. Emission intensities of dTomato (561 nm, *magenta*) and mNG (488 nm, *teal*) recorded from single endosomes using fluorescence microscopy. Emission intensities were recorded before and after photobleaching of tomato by direct exposure to 561 nm light. (**C**) FRET efficiency (E_app_) between Gα_S_ ^mNG^ and βarr1^Tom^ as a function of the acceptor-to-donor fluorescence ratio (F_Tom_/F_mNG_), measured in endosomal regions of LA-PTH-stimulated cells. The curve is a one-phase association fit; a straight line was rejected in favor of this model by an extra sum-of-squares *F* test (*F*(1, 21) = 13.15, *P* = 0.0016). Fitted plateau, 0.24 (95 % CI: 0.19 to 0.29); *R*² = 0.40. **(D)** Example of simultaneous dual-color single molecule trajectories of Gαs^mNG^ (*teal*) and βarrestin2^PAmCherry^ (*magenta*) after stimulation with 1 µM LA-PTH. **(E)** Diffusion coefficient comparison between Gαs^mNG^ (*black*) and βarr2^PAmCherry^ (*magenta*). Bar graph represents the mean of n = 278/17/26 (basal) and n=592/198/34 (LAPTH) average number of trajectories Gαs^mNG^/βarr^Pam^/number of cells for N = 3 independent experiments. ****P < 0.0001 by t-test with paired comparison. **(F** and **G)** Survival colocalization plot for random collision control (*black*), Gαs^mNG^ (*magenta*) and Gαs^mNG^ +LAPTH (t*eal*) with βarr2^PAmCherry^ (C). Half-life time of association. Parameters were obtained from the exponential decay fit of plots like C (D). Data presented as mean ± SEM from n = 8 (control), n = 9 (Gαs^mNG^/βarr^Pam^) and n = 18 cells (Gαs^mNG^/βarr^Pam^ +LAPTH). ns, non-significant with P > 0.05 by one-way ANOVA with Tukey post-hoc multiple comparison. **(H)** Time courses and integrated responses of cAMP production in MC3T3 cells after brief stimulation with 1 nM PTH, without (control, *black*) or with the endosomal Gas inhibitor (*teal*). cAMP responses are normalized to forskolin stimulation. Bars represent as mean ± S.D. (n = 32 for control; n = 18 for inhibitor). **(I)** Time courses and schematic of PTH1R internalization measured using PTH1R^SEP^ in HEK293-R cells upon PTH stimulation (mean ± S.D., n = 3).

We therefore measured E_app_ by acceptor photobleaching as a function of F_Tom_/F_NG_ in LA-PTH-stimulated cells, separately at the plasma membrane and endosomes, and compared a straight line with a one-phase association model using an extra sum-of-squares *F* test. In endosomes, the FRET efficiency increased and then plateaued as the acceptor-to-donor ratio rose (**Fig. 2C**) and the hyperbolic binding model was the preferred over the linear model (*F*(1, 21) = 13.15, *P* = 0.0016) (**Fig. 2C**). By contrast, application of the same analysis to the plasma membrane did not reject the linear model (*F*(1, 9) = 0.8383, *P* = 0.3838) as expected for proximity-dependent transfer between molecules that are not in complex (**Fig. 2C**). The saturating dependence seen only in endosomes therefore indicates a specific Gαs–βarr1 interaction confined to that compartment.

However, we did not observe a similar range of acceptor-to-donor ratios in either compartment. On endosomes, βarr1 was enriched relative to Gαs, and F_Tom_/F_NG_ reached values of up to ∼130; at the plasma membrane, the ratio did not exceed ∼6. The endosomal curve saturated above a ratio of approximately 15, which was outside the range accessible at the cell membrane. Therefore, the absence of saturable E_app_ values at the plasma membrane could reflect the lower acceptor/donor ratio detectable there, as well as the non-specificity of the Gαs-βarr interaction. Complementary single-molecule data support the latter conclusion, showing no Gαs–βarr colocalization events at the plasma membrane despite strong agonist-induced co-enrichment of the two proteins there (**Fig. 2D–G**).

Dual color single-molecule tracking of Gαs^mNG^ and βarr2^PAmCherry^ at the plasma membrane of HEK293 cells expressing the PTH1R using TIRF gave mean diffusion coefficients of 0.008 ± 0.010 µm² s⁻¹ for βarr2^mNG^ and 0.19 ± 0.13 µm² s⁻¹ for Gαs^mNG^ in unstimulated cells, a difference of approximately twenty-fold and neither was significantly altered by LA-PTH, 0.062 ± 0.07 and 0.19 ± 0.14 µm² s⁻¹, respectively (**Fig. 2D, E)**.

We further quantified the temporal colocalization between single molecules of Gαs^mNG^ and βarr2^PAmCherry^. As a negative control, we compared the colocalization time for an unrelated protein (PH lipid binding domain) and βarr2^PAmCherry^. The survival function of colocalization plot showed similar decay that our unrelated protein, as a control of random collisions. The difference in the molecular association half-life times (t_1/2_) between the two treatments was not statistically significant (**Fig. 2F, G**). Because two proteins within a single complex must diffuse as one particle, the persistence of this difference after agonist indicates that most surface Gαs does not diffuse with βarr2, consistent with the absence of specific Gα_S_–βarr interaction events at the plasma membrane. Taken together, these experiments identify a specific Gα_S_ –βarr1 interaction on endosomal membranes of agonist-stimulated cells, while no Gα_S_–βarr colocalization was detected at the plasma membrane.

### Endosomal Gα_s_ activity is required for sustained cAMP production

We next tested whether endosomal Gαs is functionally required for sustained signaling. A Gαs inhibitor targeted to endosomes, endo-Gαs-BP2 (26), markedly reduced the sustained phase of the PTH-induced cAMP response and lowered the integrated cAMP response by approximately 34% (**Fig. 2H**), without altering receptor internalization. The latter was monitored in cells stably expressing PTH1R N-terminally tagged with superecliptic pHluorin (SEP), a pH-sensitive GFP variant whose fluorescence is quenched in the acidic lumen of endosomes (27) (**Fig. 2I**). Endosomal Gαs activity is therefore required for the second cAMP wave rather than for receptor trafficking.

### Molecular simulations reveal the propensity of Gαs to dislocate away from βarr1

Because cytosolic GTP exceeds GDP by roughly tenfold, nucleotide binding is expected to activate Gαs and destabilize the PTH1R–Gα_S_–βarr1 megaplex, predicting a short-lived assembly in cells. Our live-cell FRET measurements nevertheless detect the Gα_S_–βarr interaction on endosomes, indicating that the complex persists long enough to be observed. To reconcile these observations, we performed WE-MD simulations of the conformational dynamics of the PTH1R megaplex in the absence of nucleotide. WE is an enhanced sampling strategy that, when used with MD, enables efficient exploration of conformational space beyond the reach of conventional MD (28–30), as described in the Methods and demonstrated in a recent application to class A GPCR activation (31).

We initiated the WE-MD simulations from the cryo-EM structure of the PTH1R-βarr1-Gα_S_ megaplex reported separately (25), shown in **Figure 3A**. The megaplex was embedded in a POPC (1-palmitoyl-2-oleoyl-glycero-3-phosphocholine) bilayer; LA-PTH was replaced by PTH; and the missing Gα_s_ elements were rebuilt, with engineered residues reverted to wild type, as described in the Methods. We generated more than 500 WE-MD iterations and monitored in each run the progress along two coordinates: Gα_S_ root-mean-square deviation (RMSD) relative to its original conformation, and distance between the centroids of Gα_S_ and PTH-PTH1R-βarr1 complex. The simulations yielded the ensemble of trajectories displayed in **Figure S3A**, as a function of Gα_s_ RMSD, reaching Gα_S_ RMSDs of ∼40 Å. One continuous trajectory corresponding to this partially dissociated state is highlighted in *gray* in **Fig. S3A**, illustrating the progressive increase in Gα_S_ RMSD over successive WE iterations (*ordinate*). The corresponding final conformation (**Fig. 3B**) shows an extensive displacement in the Gαs subunit, as can be seen from the structural alignment of the initial and final conformations in **Fig. 3C**. **Figure 3D** displays the time evolution of the RMSD from the original coordinates for the four components of the megaplex. Gαs is distinguished by its large RMSD increase consistent with its extensive dislocation from its original position; βarr1 shows a relatively modest conformational change (up to 10–12 Å RMSD), and the receptor and the peptide remain stable throughout (< 5 Å RMSD). Notably, the destabilization of Gαs is accompanied by an increase in TM3-TM6 distance (**Fig. 3E**), indicating the opening of the cavity where Gαs inserts upon activation. Further examination reveals that Gαs practically undergoes a rigid-body rotation away from βarr1, maintaining its overall fold, except for the reorientation of its terminal helix, which is relatively decoupled from the globular fold (**Fig. 3F** and **S3B**).

**Figure 3.**
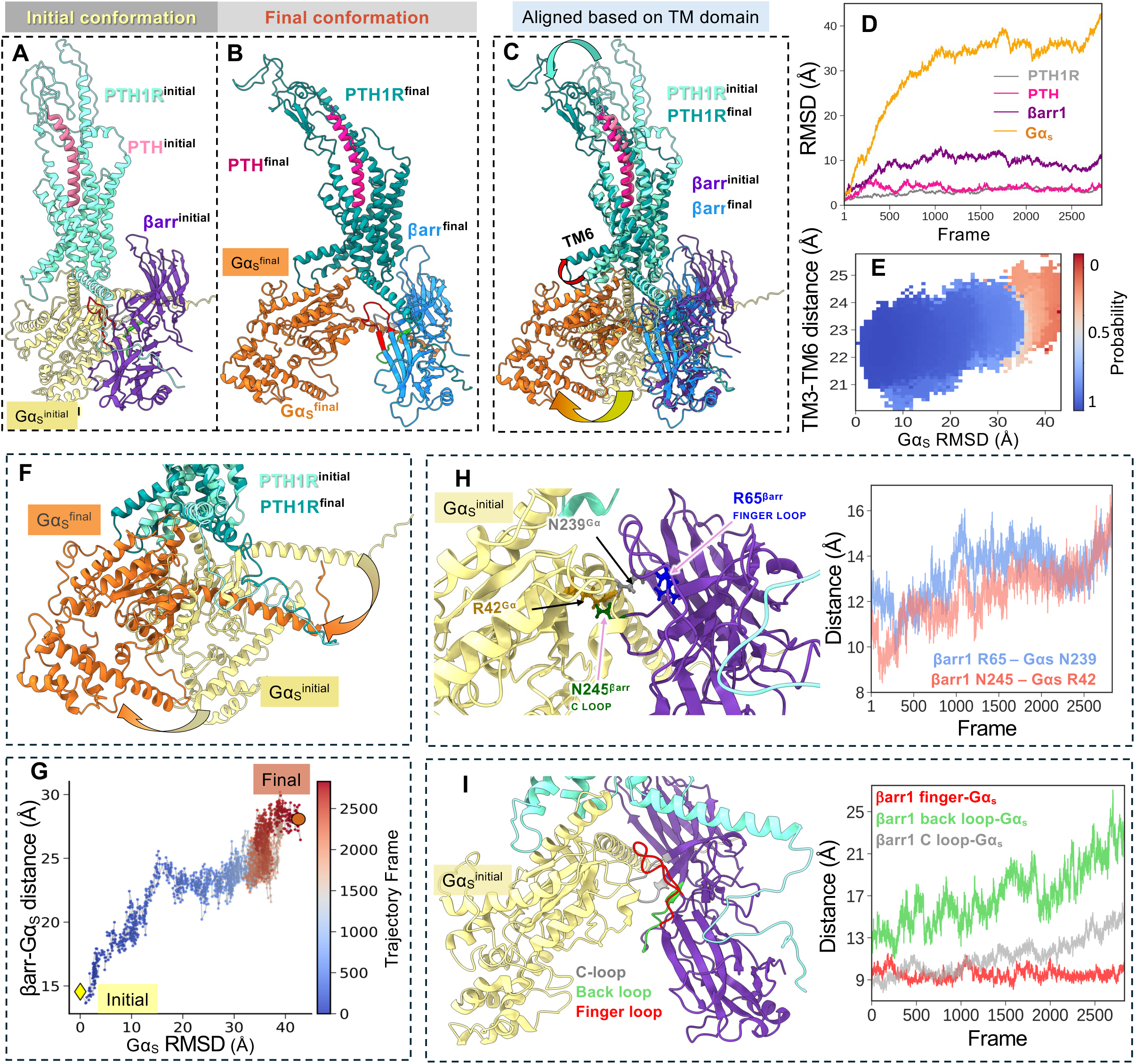
Partial dissociation of Gα_s_ from the megaplex by rigid-body reorientation away from βarr1 observed in WE-MD simulations. **(A–C)** Structure of PTH‒PTH1R‒βarr1‒Gα_S_ megaplex used to initiate WE-MD simulations (A), the final conformation observed at the end of the example trajectory highlighted in **Fig. S3A** (B), and their structural alignment upon optimal superposition of the transmembrane domain (C), illustrating the significant destabilization of Gαs binding away from βarr1. **(D)** Time evolution of the RMSDs from initial conformations in the example trajectory, plotted for PTH1R (*gray*), PTH (*magenta*), βarr1 (*purple*), and Gα_S_ (*orange*). **(E)** Evolution of TM3-TM6 distance as a function of Gα_S_ RMSD, in all the WE-MD trajectories. The color scale represents probability of occurrence, with *blue* indicating the most probable regions and *red* the least probable. **(F)** Zoomed-in comparison of the initial (*yellow*) and partially dissociated (*orange*) conformations of Gα_S_. βarr1 is not displayed for clarity. **(G**) Increase in Gα_S_ –βarr1 distance accompanying Gα_S_ destabilization. Projection of the example trajectory (*gray* in Fig. S3A) onto a 2D progress coordinate space defined by Gα_S_ RMSD (from its original position) and βarr^S341^-Gα_S_^R38^ distance. The trajectory is colored by simulation progression. **(H, I)** Interfacial interactions between Gα_S_ and βarr1 and their disruption during partial Gα_S_ dissociation. Time evolutions of two representative interfacial distance pairs, labeled in the ribbon diagram (*left*) and curves (*right*), are shown in panel **H**, and those of distances between Gαs and the finger, B-, and C-loop of βarr1, are shown in panel **I,** where the finger-, C-loop, and B-loop are colored *red*, *gray*, and *green*, respectively, in both the ribbon diagram and curves. The centroids of the following groups of residues are used as reference for the respective interactions in panel **I**: βarr1 finger loop (R65, V70, L71, F75, K77) and Gα_s_ (K233, W234, Q236, F238, N239, R280, W281); βarr1 C-loop (C242, L243, N245, T246) and Gα_s_ (F208, F222, W234, C237, F238, N239, V241); and βarr1 B-loop (R312, E313) and Gα_s_ (N66, S205).

To understand whether the extended TM6 opening accompanying Gαs reorientation is a general feature of the ensemble of trajectories, we evaluated the TM3^L306^–TM6^R404^ distances sampled during all WE-MD simulations (**Fig. 3E**). While a broad distribution of conformers is observed, the TM3-TM6 distance tends to increase with increasing Gαs RMSD from its original position, the widest cytoplasmic openings (TM3–TM6 ≈ 25 Å) occurring with the largest Gα_s_ displacement (RMSD > 35 Å). TM3–TM6 distances of 21–24 Å are those most frequently observed, but further opening of the cytoplasmic pocket is permissive, and possibly required for, Gαs dissociation.

As Gα_s_ dislocates from its binding site by a near-rigid body rotation (*curled arrow* in **Fig. 3F**), βarr1 also progressively moves away from Gαs, e.g., the distance between the centroids of residues βarr^S341^ and Gα_S_^R38^ increases from ∼15 Å to ∼28 Å (**Fig. 3G**). **Figure 3H** shows the interfacial interactions between: (i) βarr1 finger loop residue R65 and Gα_S_ N239, and (ii) βarr1 C-loop residue N245 and Gα_S_ R42 (*left panel*), the respective distances of which gradually increase during Gα_S_ dislocation (*right panel*). Additional analysis showed a gradual increase in the distance between Gα_S_ and the βarr1 back and C loops, whereas contacts with βarr1 finger loop remain minimally affected (**Fig. 3I**). The WE-MD simulations therefore demonstrated the disposition of Gα_S_ to dislocate from its binding site on the receptor, and globally reorient away from βarr1, manifested by the disruption of interactions between structural elements at the interface of Gα_S_ and βarr1.

### Simulations further show that βarr1 undergoes conformational rearrangements to stabilize its interactions with the receptor upon destabilization of Gα_s_

As noted above, Gα_S_ large-scale motions towards dissociation from the receptor are accompanied by conformational rearrangements in βarr1, the interfacial interactions between Gα_S_ and βarr1 being increasingly weakened and disrupted. Concurrently, the extracellular domain of PTH1R bends slightly along with the bound PTH, and another notable change is the movement of βarr1 closer to PTH1R to achieve tighter interaction with the receptor (**Fig. 4**). The stronger interactions are evidenced by a progressive decrease in the distance between PTH1R^helix8^ (e.g. K472) and βarr1 finger loop residues (L68-D69) (**Fig. 4A**). A similar trend was observed when evaluating the center-of-mass distance between PTH1R^ICL2^ and βarr^R331-S340^ (**Fig. 4B)**. Together, these analyses indicate a net movement of βarr1 toward PTH1R during Gα_S_ dissociation (**Fig. 4C)**. Consistent with this rearrangement, the van der Waals interaction energy between PTH1R and βarr1 becomes more favorable by about 30 kcal/mol at later stages of the simulations (**Fig. 4C**, *right panel*).

**Figure 4.**
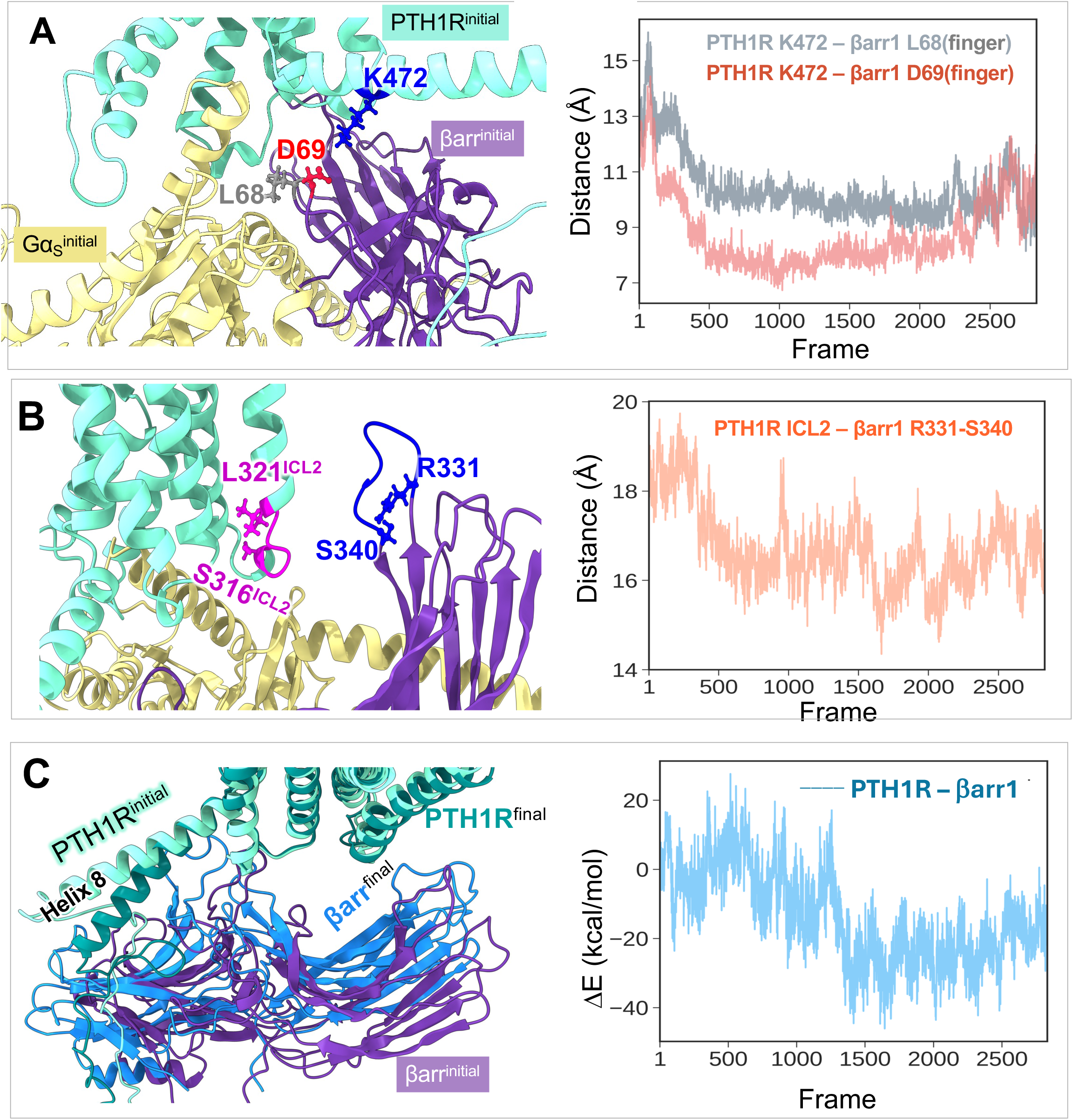
Coordinated rearrangement of PTH1R megaplex promotes closer βarr1 association with the receptor during partial Gαs dissociation. **(A-B)** Close-up view of key interactions at the PTH1R-βarr1 interface (*left*) and their time evolution (*right*), shown for interactions between PTH1R K472 and βarr1 finger residues L68 and D69 (**A**), and between PTH1R ICL2 residues and βarr1 K321 and D316 (**B**). **(C)** Zoomed-in comparison of initial and final conformations from the representative WE-MD trajectory, illustrating the movement of βarr1 closer to the receptor (Gαs omitted for clarity) (*left*); and change in van der Waals interaction energy between PTH1R and βarr1, showing the gradual strengthening of interactions as βarr1 conformational changes induce tighter contacts with the receptor (*right*).

Overall, the simulations show that Gα_S_ is poised to dissociate from the megaplex while βarr1 engages the receptor more tightly — a hand-off geometry in which loss of the G protein subunit leaves a more stably arrestin-bound receptor behind. **Supplementary Movies 1** and **2** present time-resolved views of whole megaplex dynamics (**Movie 1**) and the Gα_s_–βarr1 interfacial region (**Movie 2**).

### GTP binding promotes the disengagement of Gαs from βarr1

To test whether nucleotide binding influences the stability of the megaplex, we constructed a GTP-bound megaplex model by introducing the nucleotide into the nucleotide-binding region of Gαs (see Methods), and ran three independent 1 µs conventional MD simulations for each of the nucleotide-free and GTP-bound megaplex systems, giving 6 µs of unbiased sampling. Unlike the weighted-ensemble simulations, these MD runs report the spontaneous behavior of the megaplex with the nucleotide as the only variable, allowing direct comparison between the two conditions (with and without GTP). Comparing the two conditions revealed differences at the βarr1-Gαs interface, particularly involving the βarr1 C-loop (**Fig. 5A**), which showed increased interfacial distances in the presence of GTP. Similar but less pronounced increases in the interfacial distances were observed in the case of the βarr1 B-loop (**Fig. 5B**) and finger loop (**Fig. 5C**). Additionally, the separation between βarr1 R65 and Gαs N239 increased in the GTP-bound simulations (**Fig. 5D**). Other interfacial regions showed modest or minimal changes between the two conditions (**Fig. 5E, F**). This comparison unveiled two key features. First, the nucleotide effect is largest at the C-loop, which contacts the Switch I and Switch II surfaces in the reported cryoEM structure (25). Since Switches I and II undergo conformational change upon GTP binding, these MD simulation results show the expected allosteric effect. Second, the receptor–βarr1 interface was not correspondingly destabilized (**Fig. 5F**), indicating that nucleotide binding selectively weakens the Gα_S_–βarr1 side of the assembly rather than fully disassembling the complex.

**Figure 5.**
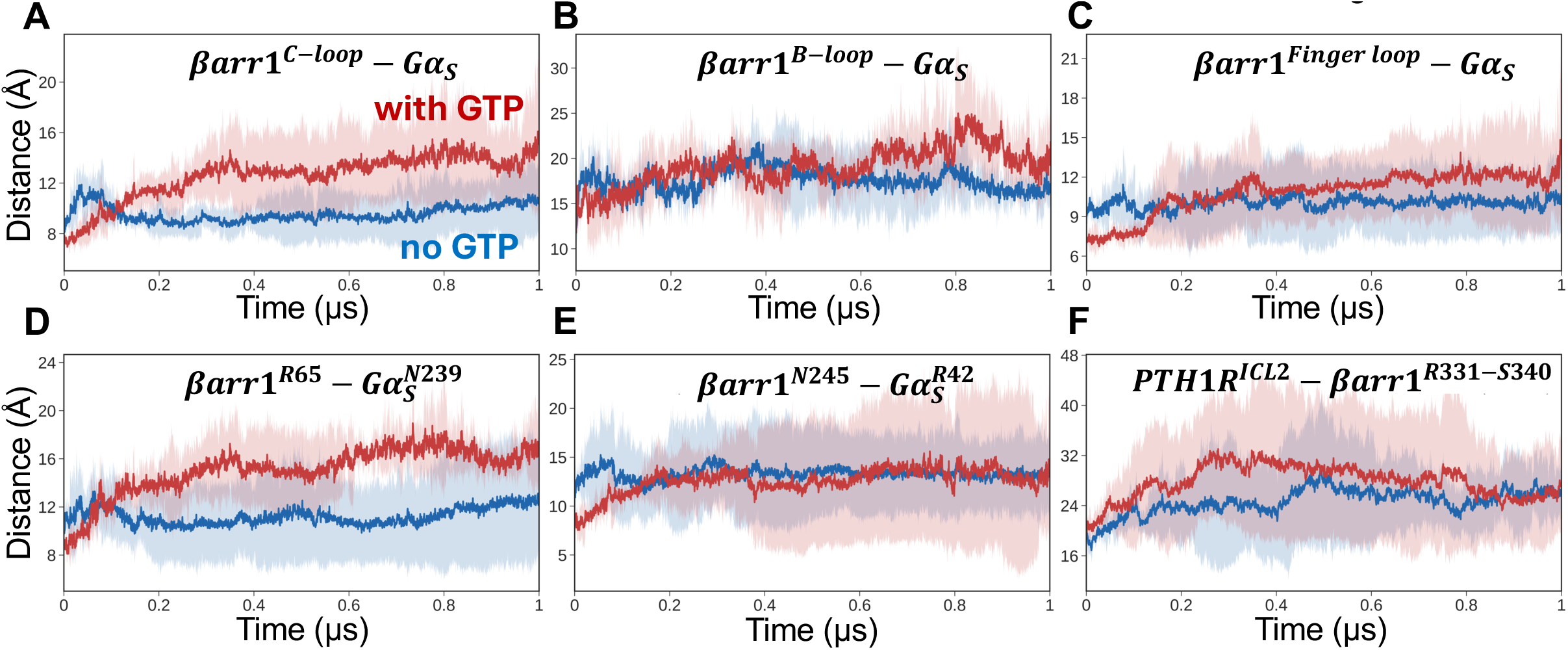
Enhancement in βarr1-Gαs distance in the presence of GTP, compared to that observed in the absence of GTP. Comparison of the PTH1R megaplex in the presence (*red*) and absence (*blue*) of GTP. Three independent 1 μs long conventional MD simulations were performed for each case, totaling 6 μs cumulative MD sampling. Solid lines show the mean across three independent runs, while shaded regions indicate standard deviation (SD). **(A–C)** Time evolution of distances between Gαs and the βarr1 C-loop (A), B-loop (B), and finger loop (C). **(D, E)** Time evolution of representative interfacial distances between βarr1 R65 and Gα_s_ N239 (D) and between βarr1 N245 and Gαs R42 (E). **(F)** Time evolution of the centroid distance between PTH1R ICL2 (S316-L321) and βarr1 (R331-S340).

In one of the three GTP-bound MD runs (MD3), the nucleotide itself was displaced from its initial binding region after ∼0.4 μs (**Fig. S4A** and **Supplementary Movie 3**). This was not observed in the other two runs; therefore, we reported it as an isolated observation. However, it is consistent with the conformational plasticity of the nucleotide-binding site in the receptor-engaged conformation. Comparison of the initial and final conformations in that trajectory revealed that the considerable displacement of the GTP relative to its initial nucleotide-binding region was accompanied by rearrangements in the α-helical domain of Gα_S_. These observations suggest that GTP-binding pocket of Gα_S_ can undergo conformational rearrangements leading to the exposure of GTP-binding and altering nucleotide accessibility. Such structural changes may be relevant to GDP/GTP exchange, although additional simulations are required to establish complete nucleotide dissociation or exchange mechanisms.

Weighted-ensemble enhanced MD simulations of the GTP-bound megaplex, performed with the same protocol as for the nucleotide-free system, were consistent with the conventional MD observations, showing comparable rigid-body displacement of Gα_S_ with somewhat greater separation at selected interfacial contacts (**Figs. S5–S7**). **Figure S7** depicts the counterparts of **Figures 3** and **4**; and **Figure S5** is the counterpart of **Figure S3**. **Figures S7** (panels **A-E)** and **S5B** again show substantial displacement of Gα_s_, primarily due to rigid-body displacement of this subunit, accompanied by a pronounced reorientation of the terminal helical region as in previous WE-MD simulations. **Supplementary Movie 4** shows the time evolution of conformational changes corresponding to the trajectory highlighted in *gray* in **Figure S5A**. GTP remained relatively close to its initial nucleotide-binding region during these rearrangements, as indicated by a comparatively small GTP RMSD of 3-4 Å after alignment of the initial and final Gα_s_ conformers (**Fig. S6A**). Comparison of example WE-MD trajectories from the GTP-free and GTP-bound megaplex reveals a comparatively steeper increase in Gαs RMSD in the presence of GTP, reaching a slightly larger displacement (∼48 Å versus ∼43 Å; **Figs. 3D** and **5D**). We also observe greater separation at selected βarr1-Gα_S_ interfacial interactions (**Fig. S7J**) and larger fluctuations in interfacial distances, suggesting that GTP binding facilitates conformational rearrangements by weakening interfacial interactions within the megaplex, favoring Gα_S_ disengagement.

Like the nucleotide-free simulations, larger Gαs displacements were associated with greater opening of the PTH1R intracellular cavity, as indicated by the 2D-scatter plot of Gα_S_ RMSD and the TM3-TM6 distance across all runs (**Fig. S7E**). Additional analysis of representative interfacial residue pairs between βarr1 and Gα_S_ revealed patterns similar to those observed in the absence of GTP (**Fig. S7H, I**).

The interactions between PTH1R-βarr1 interfacial residues became tighter but remained weaker than those observed in the absence of GTP, suggesting that the large-size fluctuations in Gαs position perturbed the association between the receptor and βarr1 (**Fig. S7J, K**), as also manifested by the fluctuations in PTH1R-βarr1 interaction energy (**Fig. S7L**). Yet, in both the GTP-free and -bound WE-MD simulations, the PTH1R-βarr1 interface became more compact during partial Gαs dissociation. The structural alignment of the initial and final PTH conformations revealed minor bending near the C-terminal region of the peptide (**Fig. S6C**). Interestingly, structural studies of LA-PTH-bound PTH1R have reported similar C-terminal rearrangements, where partial disengagement of the peptide from the receptor extracellular domain was proposed to contribute to ligand dissociation-rebinding dynamics (16, 32).

## Discussion

While this work was in preparation, a cryo-EM structure of an artificially stabilized agonist-bound PTH1R–βarr1–Gα_S_ complex was reported, together with mutational evidence that βarr1 engages Gα_S_ through a three-node interface formed by its finger, C-, and back loops (25). That study defines the interaction architecture but does not establish whether the assembly occurs in endosomes in live cells or how it resolves. The present work provides that evidence, converting a structurally defined interaction into a localized endosomal signaling metastate.

We used three imaging resolutions to determine whether the Gαs-βarr is present in endosomes. Confocal microscopy shows nucleotide-free Gαs, βarr1, and PTH-occupied receptor on the same endosomes but cannot report their arrangement on the endosomal membrane. Single-molecule TIRF and HILO microscopy at 44 nm and 96 nm resolution, respectively, separate the plasma membrane from the endosomal membrane within single cells, allowing examination of the molecular Gα_S_-βarr pair in each compartment. As expected, the agonist increased Gα_S_-βarr proximity at the plasma membrane because βarr is recruited from the cytosol to the plasma membrane, where Gαs is already resident. This response validates the proximity analysis’s sensitivity. However, at that same membrane, the complementary assays detect no interaction: no single-molecule colocalization events, and saturable FRET is detected only on endosomes, indicating that the productive Gα_S_–βarr complex assembles on the endosomal membrane. Second, the functional requirement is endosome-restricted rather than trafficking-dependent: an endosome-targeted Gαs inhibitor abolishes the sustained cAMP phase while leaving receptor internalization intact. This distinction matters because approaches that block endocytosis (e.g., dynamin inhibition, dominant-negative dynamin, or phosphorylation-deficient receptors) necessarily compare a receptor that internalizes with one that does not and therefore establish that the sustained phase requires endocytosis without localizing the responsible complex. Inhibiting Gαs only on endosomes, in cells where the receptor still internalizes normally, places the requirement in the compartment itself.

At HILO resolution, the localization reconstructions show that Gα_S_ -βarr complexes are coincident on endosomes, but they are infrequent, and most Gαs is opposed to rather than coincident with βarr-occupied endosomes. This observation indicates that the Gα_S_ -βarr complex occupies only a small fraction of the available molecules, possibly reflecting its transient state. The WE-MD simulations provide support for this conclusion. In the absence of nucleotide, the assembly is metastable rather than static: Gα_S_ escapes via a large rigid-body rotation gated by further opening of the receptor’s cytoplasmic cavity, a route with a substantial barrier that permits a lifetime long enough for detection. In this model, the complex is productive especially because it is unstable: a stable assembly would sequester Gα_S_ rather than turn it over, and sustained cAMP production from endosomes containing limited amounts of Gα_S_ would require turnover. We propose that βarr does more than sterically tolerate Gas binding; it holds the receptor in a state that can be re-engaged, which is required for the multiple rounds of Gα_S_ activation implied by sustained endosomal cAMP production (20). Our simulations also indicate which loops disengage first: contacts between Gα_s_ and the βarr1 back and C loops lengthen progressively, whereas the finger-loop contact is least affected. Within the three-node interface described structurally (25), the nodes therefore appear not to be equivalent, and the finger loop is the most persistent of the three, a prediction that interface mutagenesis alone would not reveal, since loop substitution reports the requirement for a contact rather than its order of release.

In a series of unbiased simulations, GTP consistently increased the separation between Gαs and βarr1, with the largest effect at the C-loop, which contacts the Switch I and Switch II surfaces; the receptor–βarr1 interface was comparatively unaffected. The detected asymmetry therefore implies that the complex does not simply dissociate but instead resolves specifically at the Gα_S_–βarr1 interface, consistent with the hand-off geometry observed in the nucleotide-free WE simulations, where βarr1 makes enhanced contacts with the receptor as Gαs is released. These simulations were performed without the Mg^2+^ ion, which would normally complete nucleotide coordination, so they reflect the effect of nucleotide occupancy rather than the fully activated state. In addition to the conformational plasticity of the nucleotide-binding site observed in one trajectory, these results point to a cycle in which the receptor–βarr1 complex binds nucleotide-free Gα_S_ and subsequent nucleotide binding then expels Gα_S_ in a form capable of activating adenylyl cyclase, leaving behind an arrestin-bound receptor available to capture another Gα_S_.

Pharmacologically, an endosomal PTH1R-Gαs-βarr megaplex defines a druggable target that is spatially separate from the plasma membrane. Compounds that stabilize or destabilize the Gα_S_–βarr interface, or that alter the TM6 opening required for Gα_S_ release, would be expected to tune the duration of the sustained endosomal cAMP signal without altering the acute response from the plasma membrane, a selectivity that would be attractive for disorders of PTH1R signaling and potentially beyond, given the conservation of endosomal cAMP signaling across several class A and B GPCRs.

## Materials and Methods

### Cell Culture and Transfections

Human embryonic kidney (HEK293) cells (ATCC #CRL-1573) and MC3T3-E1 cells (ATCC #CRL-2593) were cultured in DMEM (Corning #10027CV) supplemented with 10% FBS and 5% penicillin/streptomycin and maintained at 37° C in a humidified atmosphere containing 5% CO_2_. For transient expression, HEK293 were transfected with the appropriate cDNAs using Lipofectamine 3000 (Invitrogen # 11668019) and Opti-MEM (Gibco #51985091) for 24 hours before experiments.

### Peptides and Chemicals

IBMX (#I5879) was obtained from Sigma-Aldrich. Human PTH_1-34_ (#4011474) and LA-PTH were obtained from LifeTein LLC.

### Plasmids

Lifeact^mNG^ (#98877) was obtained from Addgene, and EPAC1 cAMP FRET sensor was originated in Martin J. Lohse lab (Uni. of Wuerzburg (Germany) (33). The plasmid encoding the selective endosomal Gα_S_ inhibitor, Endo-Gs-X (initially named Gα_S_-BP2), was developed in Mikel Garcia-Marcos lab (Boston Uni.) (34).

### cAMP recordings in single cells

Cell culture reagents were obtained from Corning (CellGro). Human embryonic kidney cells (HEK293; ATCC, Georgetown, DC) stably expressing the recombinant human HA-tagged PTH1R (HEK-293R) were grown in DMEM supplemented with 5 % FBS and 1 % penicillin/streptomycin, at 37 °C in a humidified atmosphere containing 5 % CO_2_. For transient expression, cells were seeded in six-well plates and cultured for 24 h prior transfection with the appropriate cDNAs using Lipofectamine 3000 (Life Technologies) for 24 h. Measurements of cAMP production were performed using single-cell FRET-based assays. In brief, cells were transiently transfected with the Epac1-CFP/YFP cAMP sensor. Then, cells were plated on poly-D-lysine-coated glass coverslips, mounted in Attofluor cell chambers (Life Technologies), and maintained in HEPES buffer containing 150 mM NaCl, 20 mM HEPES, 2.5 mM KCl, 1 mM CaCl_2_, and 0.1 % BSA, pH 7.40. Imaging was performed on a Nikon Ti-E microscope with a 40X oil-immersion Plan Apo objective (N.A 1.30) and a motorized stage (Nikon Corporation). For excitation, we used a mercury lamp to stimulate CFP and YFP, with fluorescence emissions captured through 480 ± 20 nm (CFP) and 535 ± 15 nm (YFP) filters. A LUCAS EMCCD camera (Andor Technology) and DualView 2 (Photometrics) with a 505 nm dichroic long-pass filter were used for simultaneous collection. Cells were perfused with either buffer, ligand or Forskolin, as indicated by the horizontal bar in the plots. We recorded fluorescence data from single cells using Nikon Element Software (Nikon Corporation). We calculated the FRET ratio (YFP/CFP) for each cell, with corrections for background, bleed-through, and photobleaching as described previously (35).

### FRET experiments

Photobleaching assays in endosomes were performed as previously described (17). Cells plated on coverslips were mounted in Attofluor cell chambers (Life Technologies) and incubated with HEPES buffer containing 150 mM NaCl, 10 mM HEPES, 2.5 mM KCl and 0.2 mM CaCl_2_, 0.1% BSA, pH 7.40, were transferred on the Nikon Ti-E microscope (Nikon) equipped with a Z-driven piezo motor. Imaging was acquired using a Nikon A1 confocal unit, through a 60 × N.A. = 1.45 objective (Nikon). Fluorescent proteins or peptides containing, GFP, mNG, Tomato, mCherry, or tetramethylrhodamine (TMR) were excited with 488-nm (GFP, mNG), or 561-nm (Tomato or TMR) lasers (Melles Griot). Data acquisitions were done using Nikon Element Software (Nikon Corporation). For FRET by acceptor photobleaching, HEK293 cells stably expressing the receptor were transfected with βarr1^dTomato^ and Gαs^mNG^ (wild-type full-length Gαs). The emission intensity of mNeonGreen was first recorded at 488-nm excitation (mNG^before^), followed by direct illumination of dTomato at 561 nm for 30-60 s. Subsequently, the emission intensity of mNeonGreen was recorded again (mNG^after^). FRET efficiency was calculated according to equation (1) (17): FRET efficiency = 1 – (mNG^before^/mNG^after^).

### Receptor internalization

We imaged receptor internalization using a Nikon A1 confocal microscope. Briefly, HEK293 cells stably expressing a pH-sensitive GFP variant, superecliptic pHluorin, inserted in the N-terminal domain of the human PTH1R, PTH1R-SEP were seeded on glass coverslips coated with poly-d-lysine (Sigma-Aldrich, P6407) for 24 hours. Experiments were carried out at 37 °C in HEPES buffer used for cAMP experiments. Cells were stimulated by 100 nM ligand for 1 min and images were acquired every 30 s. The change in fluorescence ΔF = F_0_ –F was normalized as with ΔF/F_0_, with F_0_ initial fluorescence at t = 0 min and in the absence of the ligand, and F, fluorescence at t > 0 min. The initial fluorescence was set to 100 %.

### Single-molecule localization microscopy (SMLM)

HEK293 cells were plated in 35 mm tissue culture dishes with 20 mm number 1.5 cover glass apertures (CellVis) previously coated with poly-D-lysine for 30 min at 37°C. Cells were transfected 24 hours prior the experiment using lipofectamine 3000. Transfection was performed using 50,800 and 500 ng of Gαs^mNG^ (triple expression Gαs^mNG^, Gβγ), βarr2^PAmCherry^ and PTH1R-HA plasmids respectively. For the experiment, cells were placed in 2 ml of pre-warmed (37°C) FluoroBrite DMEM (Life Technologies A1896702), supplemented with 25 mM HEPES (pH 7.4) and 5% heat-inactivated fetal bovine serum. Dual-color single-molecule time-lapses were recorded on a Nikon TiE motorized inverted microscope equipped with a Cairn OptoSplit II system, a TIRF illuminator, a 100X, 1.45 NA plan-apochromatic oil-immersion objective, and a Nikon LUN-F laser 405/488/561/640. Photograms were captured on a Prime 95B camera (Teledyne photometrics). The laser angle for TIRF and HILO configurations were calibrated with the expression of Lyn11^iRFP713^ and EEA1^GFP^, as plasma membrane and endosome markers simultaneously.

Super-resolution images were reconstructed using the ThunderSTORM plugin in ImageJ. Images were acquired at 20 Hz (50 ms per frame) for at least 1,200 frames. Camera parameters were set using a pixel size of 74 nm, an offset of 0.8, a conversion factor of 1.6 photons per analog-to-digital unit (ADU). Molecular localizations were detected using a wavelet filter based on a B-spline function (scale = 2.0; order = 3), followed by local-maximum detection with 8-neighborhood connectivity and a manual detection threshold of 3. Subpixel localization was performed using an integrated Gaussian point-spread function (PSF) model with an initial sigma of 1.6 pixels and a fitting radius of 3 pixels, using a weighted least-squares fitting method. Super-resolution images were rendered using the averaged shifted histograms (ASH) method. The analysis of the Nearest-neighbor distance (NND) was determined from the x–y coordinates of the molecular localizations identified by ThunderSTORM. Then, the closest Euclidean distance between each molecular localization of Gαs^mNG^ and its neighboring βarrs2^PAmCherry^ localization was measured. Frequency histograms were constructed using 5-nm bins, and a single Gaussian function was fitted to determine the mean distance between molecules per cell. Fits with R² < 0.5 were excluded from the analysis. The spatial resolution of the images was estimated using LifeAct^mNG^ cells imaged under the same experimental conditions used for the experiments. Two independently recordings of the same field were acquired and analyzed in ImageJ using the Fourier Ring Correlation (FRC) plugin. This analysis yielded a resolution limit of 46 ±0.002 nm for TIRF and 94 ±0.004 nm for HILO (**Fig. S2D**). For diffusion coefficient (D) measurements on TIRF mode, curves of mean-square displacement (MSD) were produced from trajectories longer than 16 frames, only the first 4-time intervals were used to fit the equation *MSD*_1−4_ = 4*Dt^α^* . Colocalization by single-molecule analysis was performed as the detection of molecules in channels 1 and 2 within 80 nm of each other in the same frame. The coordinates of trajectories in channel 1 were compared with those in channel 2 to identify colocalization events across adjacent frames. Only trajectories longer than 0.2 s were included in the analysis. Single-frame colocalization events were excluded. Interruptions in coincidental detection of up to two consecutive frames were permitted to maintain continuity of colocalization events. To generate colocalization survival plots (1 − CDF), the cumulative distribution function (CDF) of colocalization event durations of Gαs^mNG^ with respect to βarrestin2^PAmCherry^ was calculated for each cell. The resulting survival distributions were fitted with a monoexponential decay function to determine the colocalization half lifetime. Fits were performed using at least eight colocalization events. For cells with fewer than eight events, the durations were pooled across cells to reach the minimum number of events required for fitting.

### Construction of a complete structural model for PTH-PTH1R–βarr1–Gαs complex

We constructed a complete structural model for PTH-PTH1R‒βarr1‒Gα_s_ complex using the cryo-EM structure resolved for LA-PTH–PTH1R–βarr1–miniGαs. We mutated LA-PTH into PTH from S1 to F34, and constructed the missing residues of PTH1R (R58-G105, G247-A275, and A487-P520) and Gα_s_ (M1-Y37, G67-T204, and N254-T263) using AlphaFold2 (36) and PyMOL Molecular Graphics (Version 2.1, Schrödinger, LLC.). The completed model is composed of PTH (S1-F34), PTH1R (D30-P520), Gα_s_ (M1-L394), and βarr1 (T6-K357) molecules, with all mutated residues in the cryo-EM structure converted into their wt counterpart. Nine PTH1R residues were phosphorylated as pY320, pT387, pT392, pS489, pS491, pS492, pS493, pS495, and pS519. To construct the GTP-bound megaplex, we used the previously prepared nucleotide-free megaplex model and introduced GTP into the nucleotide-binding region of Gαs using the cryo-EM resolved GTP-bound dopamine D1 receptor-miniGαs complex (PDB ID: 7F23) as structural template. Specifically, the Gαs component (chain A) of the structure with PDB id 7F23 was structurally aligned with Gαs in the reconstructed PTH1R megaplex, and the coordinates of the bound GTP molecule were transferred to the corresponding nucleotide-binding region. The remaining components of PDB 7F23 were removed, retaining the original PTH–PTH1R–βarr1–Gαs assembly and the transferred GTP molecule.

Both nucleotide-free and GTP-bound megaplex models were independently embedded in a POPC lipid bilayer and solvated in 0.15 M NaCl aqueous environment using the CHARMM-GUI (37, 38) Bilayer Builder. The resulting system contained approximately 301,080 atoms. The all-atom CHARMM36m (39) force field was used for proteins, C36 (40) for lipids, and TIP3P (41) for water molecules.

### Energy minimization and equilibration

Prior to performing WE-MD simulations, both nucleotide-free and GTP-bound megaplex systems were subjected to the same series of energy minimization and equilibration steps, before production simulations. Initial energy minimization was performed without restraints for a total of 2,000 cycles, including 500 steps of steepest descent and 1,500 steps of conjugate gradual minimization. Subsequently, the system was equilibrated in three stages. First, a 20 ps restrained equilibration was performed in the NVT ensemble using a Langevin thermostat at 298 K, with harmonic positional restraints (of 1.0 kcal.mol^-1^/Å^2^) applied to all heavy atoms. This was followed by a 1 ns restrained equilibration in the NPT ensemble at 298 K and 1 atm using a Langevin thermostat and Monte Carlo barostat, maintaining the same positional restraints. Finally, a 1 ns unrestrained NPT equilibration was carried out under identical T and P. Periodic boundary conditions were applied throughout, with a 10 Å cutoff for nonbonded interactions. The fully equilibrated structures were used as the initial state for subsequent conventional MD and WEMD simulations. All minimization and equilibration steps were performed using Amber 24. To ensure compatibility, CHARMM-GUI generated input files were converted to Amber compatible force fields formats using *ParmEd* (*42*).

### Conventional MD simulations

To characterize megaplex conformational dynamics under different nucleotide conditions, three independent 1 μs long conventional MD simulations were performed for each of the nucleotide-free and GTP-bound systems, resulting in an aggregate simulation time of 6 μs. Production simulations were conducted in the NPT ensemble at 298 K and 1 atm using a Langevin thermostat with a collision frequency of 1 ps^-1^ and a Monte Carlo barostat with a pressure relaxation time of 1 ps. A 2 fs integration time step was employed, with bonds involving hydrogen atoms constrained using the SHAKE algorithm. Periodic boundary conditions were applied, and a 10 Å cutoff was used for nonbonded interactions. No positional restraints were applied during production simulations. Trajectory coordinates were recorded every 5,000 integration steps, corresponding to a 10 ps sampling interval.

### WE-MD simulations

WEMD simulations were performed using WESTPA 2.0 (43). For primary nucleotide-free and GTP-bound megaplex simulation, a 2D progress coordinate was defined using Gα_s_ RMSD and the distance between the mass centers of Gα_s_ and PTH1R-PTH-βarr1 complex. The progress coordinate space was divided into a 5 × 5 bin grid (25 total bins) using the minimal adaptive binning scheme (44). Five walkers were targeted per occupied bin, with a trajectory propagation interval (τ) of 50 ps per WE iteration. Trajectories are evaluated after each propagation interval followed by the evaluation of each trajectory using cpptraj (45) to possibly split or merge them, keeping track of their fractional weights, depending on the bins they have reached (as described in (31)). A total of 567 WE iterations were performed for both GTP-free and GTP-bound megaplex WE-MD simulations. The total weight of the generated ensemble of trajectories per iteration remains unchanged and equal to unity. Structure and trajectory visualizations were performed using ChimeraX (46) and VMD (47).

### Statistical analyses

Data were analyzed using one-way ANOVA, two-way ANOVA with a Fisher’s LSD test multiple comparison or student’s t-test as indicated in each figure legend. P values < 0.05 were considered statistically significant (\**P* < 0.05; \*\**P* < 0.01; \*\*\**P* < 0.001, \*\*\*\**P* < 0.0001). In the single-cell experiments reported here, individual cells (captured in the same dish during the same imaging session) are considered biological replicates because cell culture is asynchronous. Therefore, each measured cell is considered an independent biological measurement. These measurements were averaged across n cells from N separate experiments. Statistical analyses were performed using GraphPad Prism version 10 (GraphPad Software, Inc., La Jolla, United States).

## Graphics

Schematics in Figure 1A was created in BioRender. Vilardaga, J. (2026) https://BioRender.com/e5e0685.

## Materials and Data Availability

The source data will be deposited and made publicly available as of the publication date. Any additional information required to reanalyze the data reported in this paper and the materials will be available from the lead contacts upon request.

## Acknowledgements

We thank Drs H. Eric Xu and Li-Hua Zhao for sharing their cryoEM structure of LAPTH-PTH1R-miniGαs-βarr1 complex, determined in their laboratory and reported separately, which made the simulations in this work possible. We thank Isabelle Shimmer for technical assistance, and Dr. Jiyoung Lee for modeling the unresolved segments of the megaplex.

## Author contributions

Conceptualization: J.-P.V.

Methodology: J.P., S.S., J.-P.V., I.B.

Investigation: J.P., S.S., K.A.P

Analysis: J.P., S.S., K.A.P., J.-P.V., I.B

Funding acquisition: J.-P.V., I.B.

Project administration: J.-P.V., I.B.

Supervision: J.-P.V., I.B.

Writing – original draft: J.-P.V., I.B.

Writing – review & editing: J.P., S.S., K.A.P., J.-P.V., I.B

