## Supplementary Figures for "A metastable Gα_S_/β-arrestin complex sustains endosomal PTH1R signaling"

**This PDF file includes:**

Supplementary Figures S1–S7

Supplementary Movies 1–4

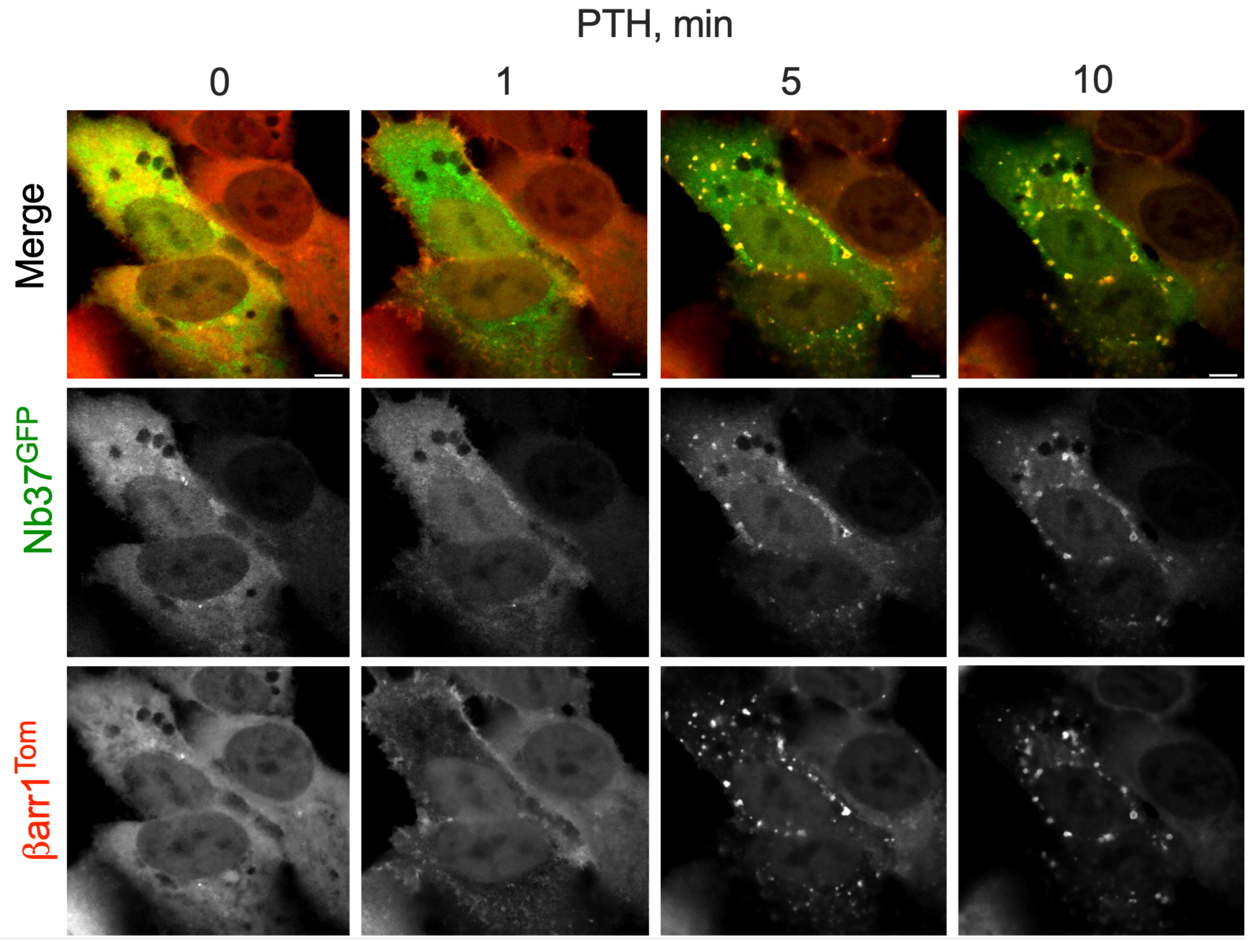


**Figure S1. Time course of confocal images related to Fig. 1A.** Confocal imaging of the distribution of $\beta arr1^{Tom}$ and $Nb37^{GFP}$ in HEK293 cells expressing PTH1R and stimulated by 10 nM PTH. The horizontal bar represents 10 µm.

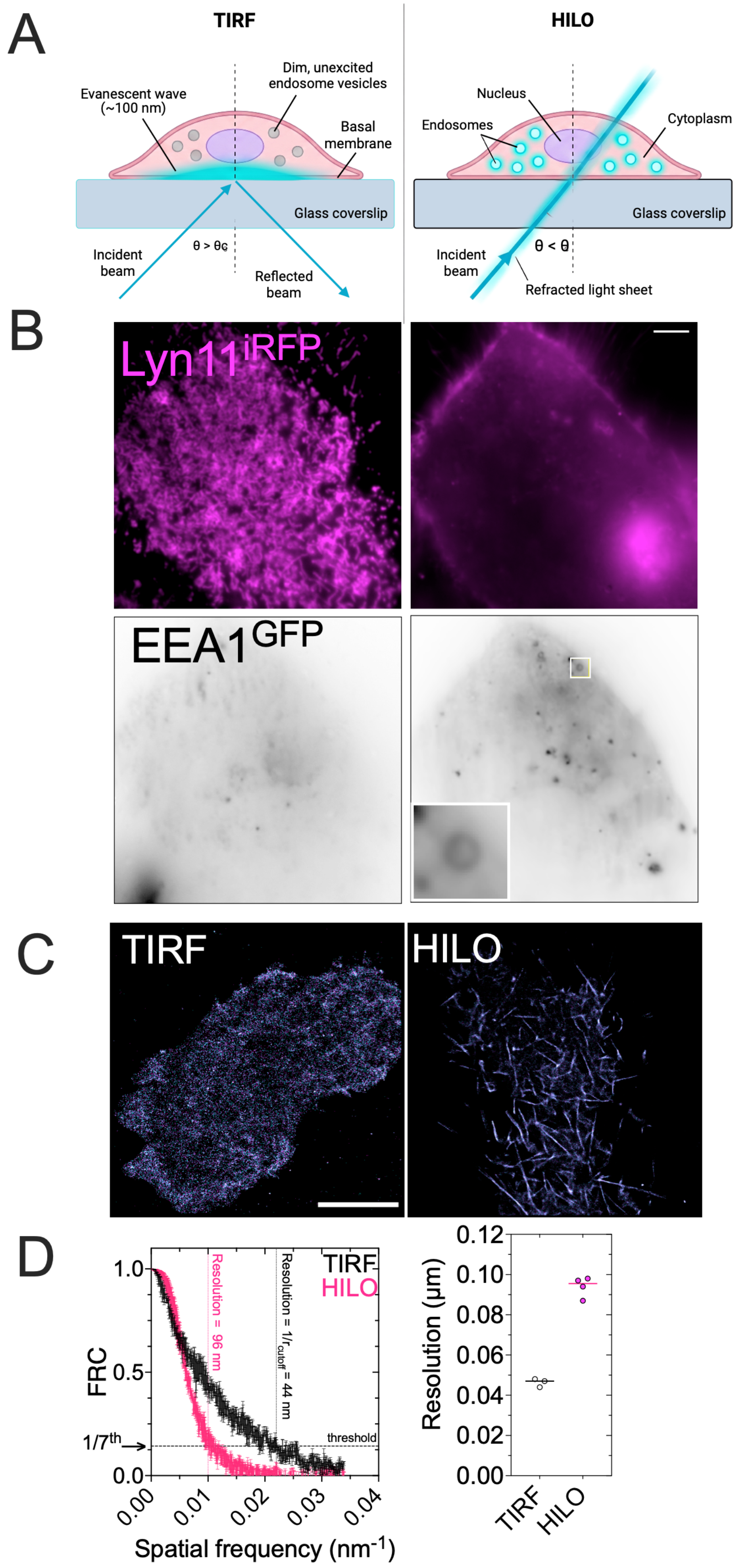

A
TIRF
HILO
Evanescent wave (~100 nm)
Dim, unexcited endosome vesicles
Basal membrane
Glass coverslip
Incident beam
θ > θc
Reflected beam
Nucleus
Endosomes
Cytoplasm
θ < θ
Refracted light sheet
B
Lyn11iRFP
EEA1GFP
C
TIRF
HILO
D
TIRF
HILO
Resolution = 96 nm
Resolution = 1/rcutoff = 44 nm
threshold
1/7th
FRC
1.0
0.5
0.0
0.00
0.01
0.02
0.03
0.04
Spatial frequency (nm-1)
Resolution (µm)
0.12
0.10
0.08
0.06
0.04
0.02
0.00
TIRF
HILO

**Figure S2. Resolution and location precision in TIRF and HILO.**

**(A)** Schematic comparing TIRF and HILO illumination.

**(B)** TIRF and HILO enable optical separation of the two compartments: the plasma membrane (TIRF) and endosomes (HILO). Representative images of a HEK293 cell expressing the $EEA1^{GFP}$ (*inverted grayscale*) acquired under TIRF (*top*) and HILO (*bottom*) illumination. Under TIRF the plasma membrane is in focus and endosomes are undetected; under HILO, EEA1GFP endosomes are resolved (*inset*) while the plasma membrane is excluded. The Lookup Table (LUT) was adjusted manually to compensate for differences in signal intensity between TIRF and HILO imaging. Scale bar, 5 µm.

**(C)** TIRF and HILO images of $Lifeact^{mNG}$ reconstruction from 3 independent recordings on the same cell, registered and analyzed by Fourier ring correlation (FRC) to establish the spatial resolution between both images.

(**D**) Spatial correlation plot used to determine the resolution of the imaging setup. Here, a 1/7 cut-off is used to establish the minimum correspondence between the 2 images. The ratio of effective pixel size (nm) to spatial frequency ($f_c$) at the cut-off indicates the system's spatial resolution.

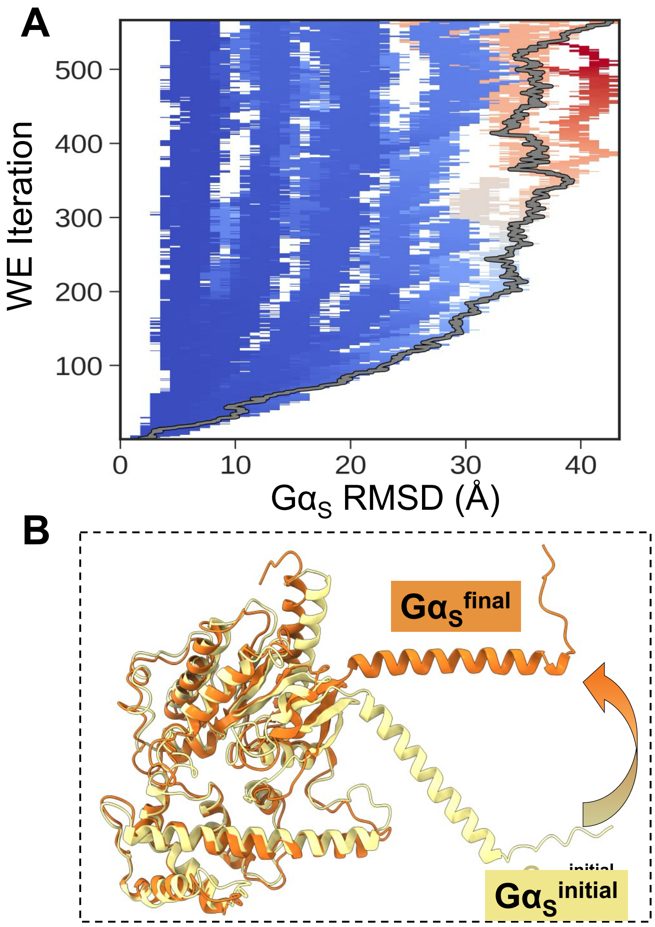


**Figure S3. Large-scale movements of $G\alpha_s$ observed in WE-MD simulations of the Megaplex**

**(A)** Ensemble of WE-MD trajectories along the progress coordinate (x-axis) defined by the $G\alpha_S$ RMSD relative to the initial conformation. The y-axis indicates the number of WE iterations, showing that RMSD values gradually increase. The curves are color-coded by their probability density (*blue*, most probable; *red*, least probable). An example trajectory reaching a $G\alpha_s$ RMSD of ~ 43 Å is highlighted in *gray*. **Supplementary Movies 1 & 2** present the time-resolved view of the highlighted pathway.

**(C)** Illustration of the rigid-body rotation of $G\alpha_S$ by structural alignment of the initial and final conformations. Note that the fold remains largely unchanged, except for the terminal helix.

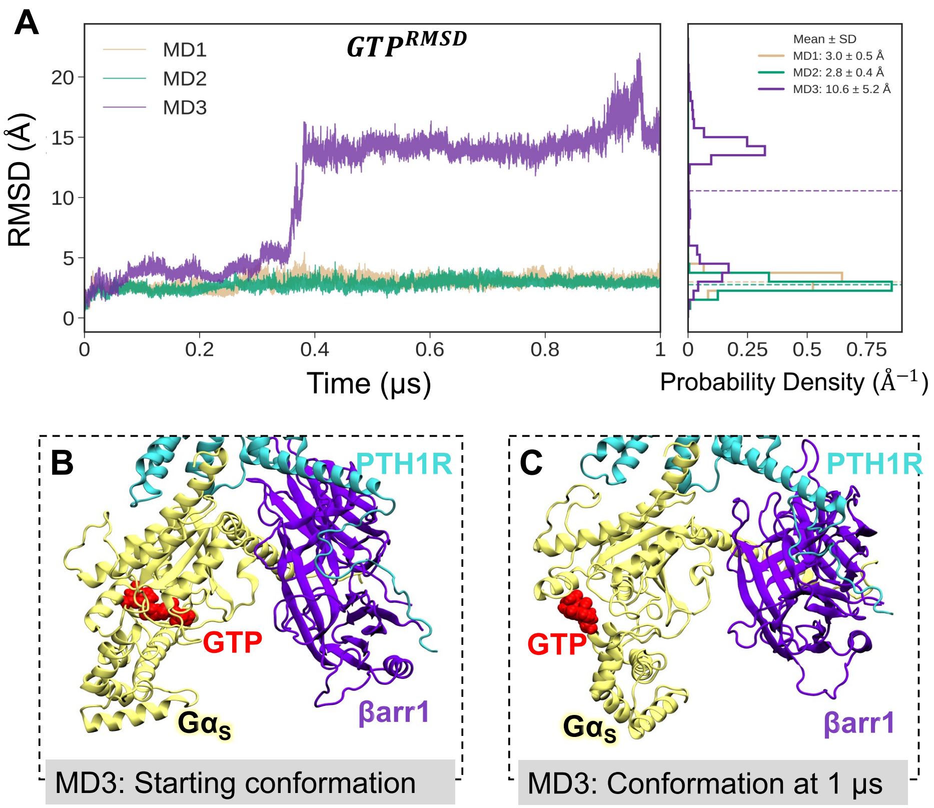


**Figure S4. Substantial displacement of GTP from its initial nucleotide-binding site during a conventional MD simulation of the PTH1R megaplex dynamics.**

**(A)** Time evolution of GTP RMSD following structural alignment of Gα$_S$, without additional fitting of GTP. While GTP remains relatively close to its initial position in MD1 and MD2, MD3 exhibits a pronounced increase in GTP RMSD around 0.4 μs, indicating substantial displacement of GTP relative to its initial configuration within Gα$_S$. Corresponding probability-density distributions are shown alongside each RMSD profile, with mean ± standard deviation (SD) calculated over the sampled trajectory frames.

**(B, C)** Initial **(**B**)** and final **(**C**)** conformations observed in run MD3, illustrating GTP dislocation from its binding site in Gα$_S$. PTH1R, βarr1, Gα$_S$, and GTP are shown in *cyan*, *purple*, *yellow*, and *red*, respectively. Comparing the two conformations reveals GTP dissociation from its initial nucleotide-binding site, consistent with the elevated GTP RMSD observed in panel A. **Supplementary Movie 3** shows the time evolution of this transition, along with the corresponding change in GTP RMSD.

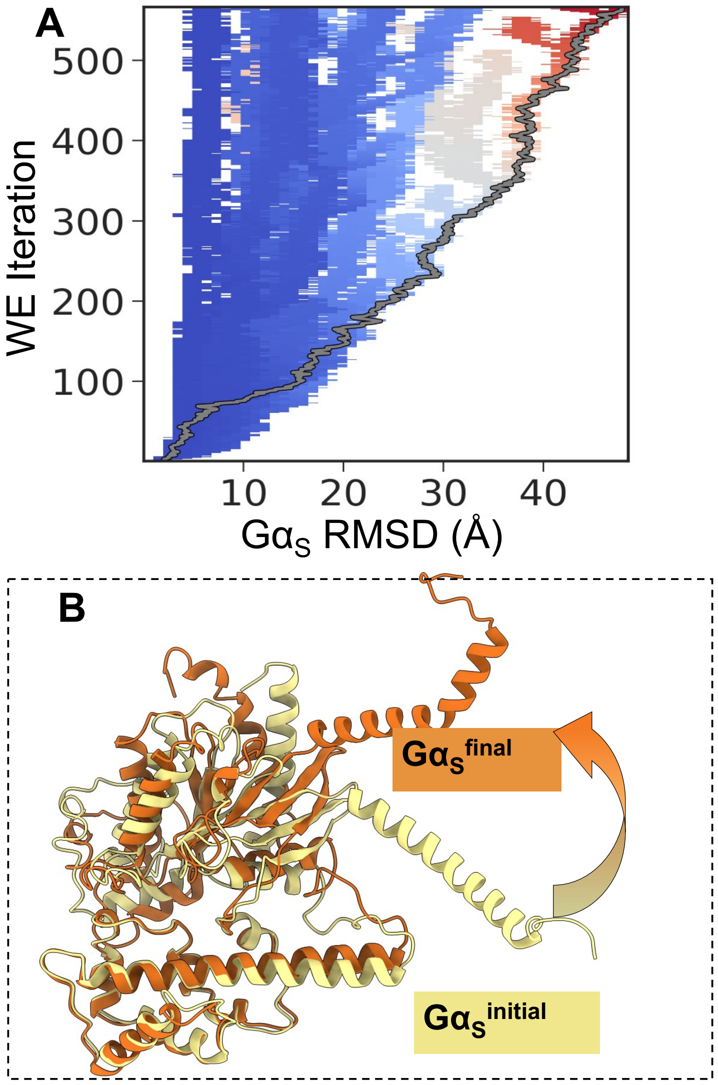


**Figure S5. Large-scale Gαs displacement and conformational rearrangement of the GTP-bound megaplex during WEM-D simulations.**

**(A)** Ensemble of WE-MD trajectories along the progress coordinate (x-axis) defined by the $G\alpha_S$ RMSD relative to the initial conformation. The y-axis indicates the number of WE iterations, showing that RMSD values gradually increase. The curves are color-coded by their probability density (*blue*, most probable; *red*, least probable). An example trajectory reaching a $G\alpha_s$ RMSD of ~ 48 Å is highlighted in *gray*. **Supplementary Movie 4** presents the time-resolved view of the highlighted pathway.

**(B)** Superposition of the initial (*yellow*) and final (*orange*) $G\alpha_S$ conformations, following independent alignment of Gαs. Apart from the terminal alpha helix, most protein regions show substantial overlap, suggesting preservation of the native fold during partial dissociation. GTP is not shown for this alignment.

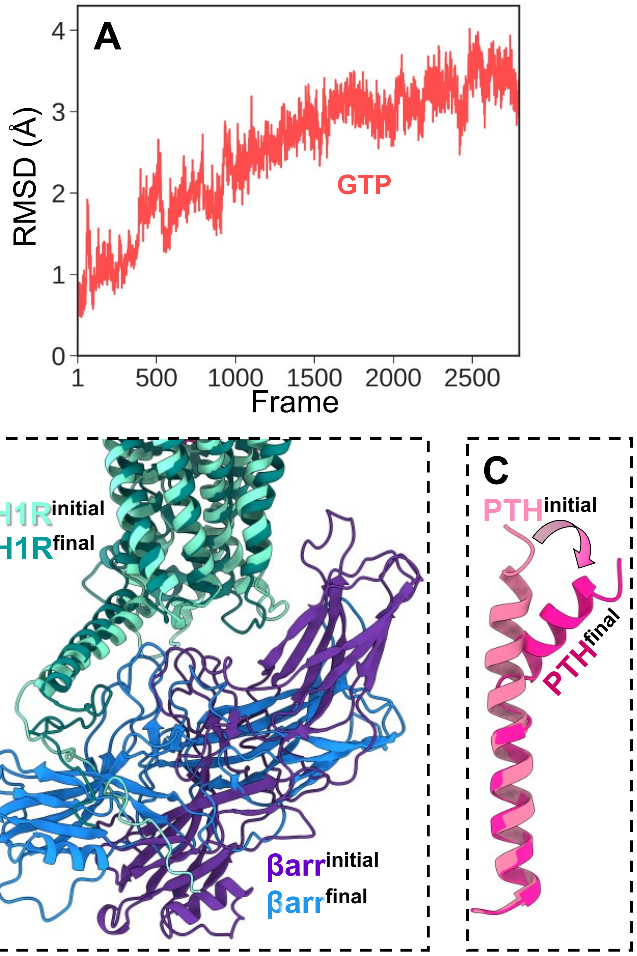


**Figure S6. Structural and interfacial rearrangements within the GTP-bound megaplex during WE-MD simulations.**

**(A)** Evolution of GTP RMSD from the example WE-MD trajectory. RMSD was calculated without additional fitting following structural alignment of $G\alpha_S$.

**(B)** Aligned initial and WE-MD-simulated final conformation (from the trajectory highlighted in Fig. 4A) of the GTP-bound megaplex to highlight the rearrangements at the PTH1R-βarr1 interface. $G\alpha_s$ is not shown for clarity in visualization purposes.

**(C)** Superposition of the initial (*light pink*) and final (*magenta*) PTH conformations, following independent alignment of PTH. A localized C-terminal bending of PTH was observed along the represented pathway.

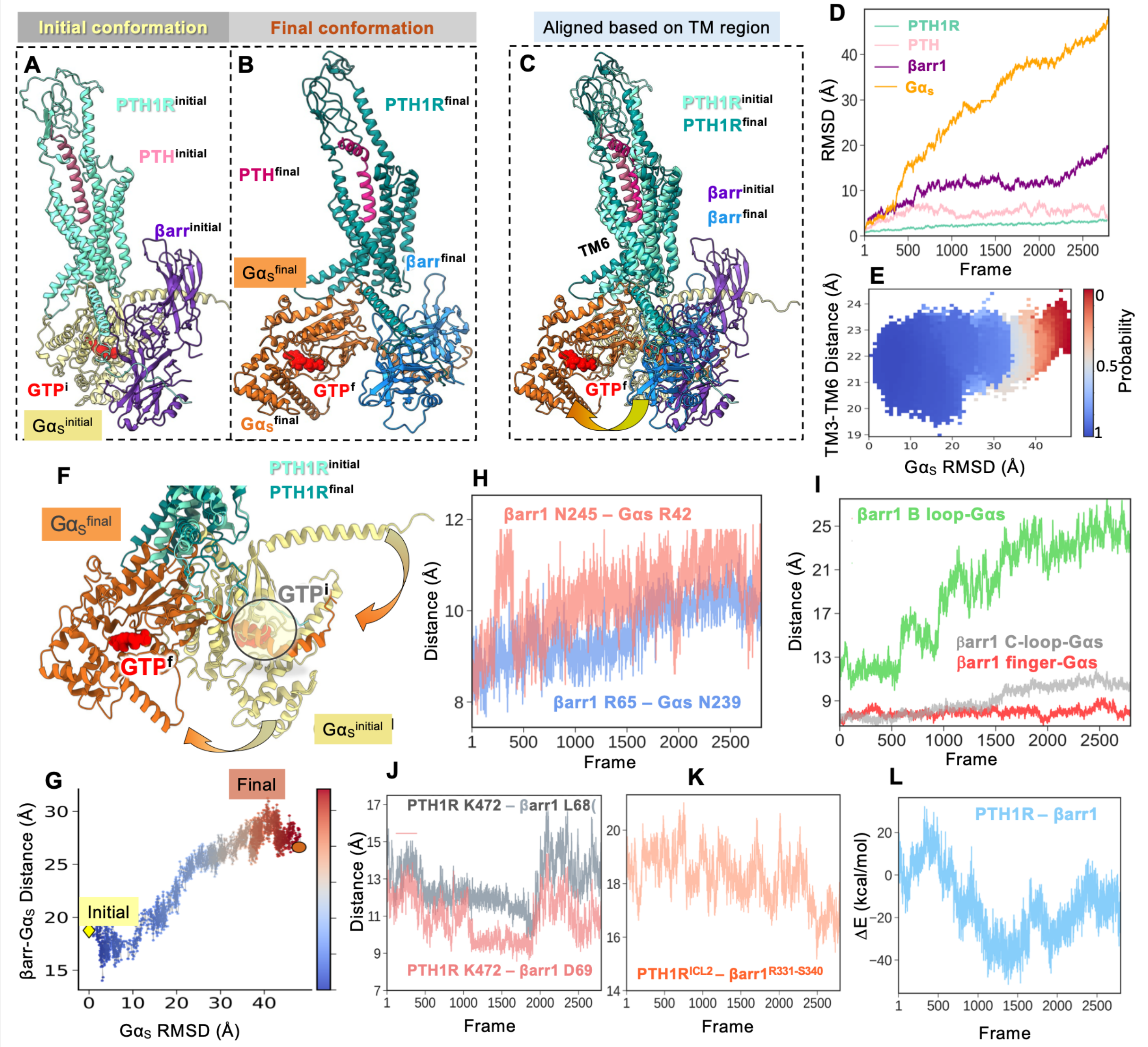


**Figure S7. Results from WE-MD simulations of GTP-containing PTH1R megaplex.**

**(A–G)** Counterparts of panels A-G of Figure 3, in the presence of GTP.

**(H, I)** Same as Figure 3H and 3I, *right* panels.

**(J–L)** Counterparts of the *right* plots in the respective panels A–C of Figure 4.

### Supplementary Movies

**Description of supplementary movies illustrating conformational rearrangements within the PTH1R megaplex during conventional MD and WE-MD simulations.**

| Movie | Movie title |
|---|---|
| Supplementary Movie 1 | G$\alpha_S$ displacement and megaplex rearrangement during GTP-free WE-MD simulation |
| Supplementary Movie 2 | Zoomed-in view of the βarr1-G$\alpha_S$ interface during partial G$\alpha_S$ dissociation |
| Supplementary Movie 3 | GTP displacement and G$\alpha_S$ $\alpha$-helical domain rearrangement during conventional MD simulations |
| Supplementary Movie 4 | G$\alpha_S$ displacement and megaplex rearrangement during GTP-bound WE-MD simulation |
